# Multi-omic profiling of the developing human cerebral cortex at the single cell level

**DOI:** 10.1101/2022.10.14.512250

**Authors:** Kaiyi Zhu, Jaroslav Bendl, Samir Rahman, James M. Vicari, Claire Coleman, Tereza Clarence, Ovaun Latouche, Nadejda M. Tsankova, Aiqun Li, Kristen J. Brennand, Donghoon Lee, Guo-Cheng Yuan, John F. Fullard, Panos Roussos

## Abstract

The cellular complexity of the human brain is established via dynamic changes in gene expression throughout development that is mediated, in part, by the spatiotemporal activity of cis-regulatory elements. We simultaneously profiled gene expression and chromatin accessibility in 45,549 cortical nuclei across 6 broad developmental time-points from fetus to adult. We identified cell-type specific domains in which chromatin accessibility is highly correlated with gene expression. Differentiation pseudotime trajectory analysis indicates that chromatin accessibility at cis-regulatory elements precedes transcription and that dynamic changes in chromatin structure play a critical role in neuronal lineage commitment. In addition, we mapped cell-type and temporally specific genetic loci implicated in neuropsychiatric traits, including schizophrenia and bipolar disorder. Together, our results describe the complex regulation of cell composition at critical stages in lineage determination, serve as a developmental blueprint of the human brain and shed light on the impact of spatiotemporal alterations in gene expression on neuropsychiatric disease.

**One-Sentence Summary:** Simultaneous profiling of gene expression and chromatin accessibility in single nuclei from 6 developmental time-points sheds light on cell fate determination in the human cerebral cortex and on the molecular basis of neuropsychiatric disease.

## Main text

Human brain development starts during the early stages of embryogenesis and extends postnatally through infancy, childhood, adolescence, and young adulthood (*1, 2*). To produce distinct circuits in the human cortex, neurons are born in an immature state and undergo a variety of molecular and morphological changes as they differentiate, migrate, and establish synaptic networks. Environmental and genetic risk factors can disrupt these highly orchestrated developmental processes, potentially leading to neuropsychiatric disease (*3, 4*). Given the variable age of onset of different neurodevelopmental disorders, it is critical to examine the effect of risk factors across the full spectrum of human brain development.

The developmental transition of cell lineages is highly orchestrated by dynamic changes in gene expression, mediated in part by spatiotemporal patterns of transcription factor (TF) binding to *cis*-regulatory DNA elements (*5–9*). Single-cell transcriptome analysis has expanded our knowledge of cellular diversity and the molecular changes that occur during differentiation, migration, and synaptic network formation in the human cortex (*9–13*). Recently, simultaneous multi-omic (gene expression and chromatin accessibility) single cell profiling has emerged as a means to decipher how combinations of TFs drive gene expression programs and to infer cell lineage transitions during development (*14*). Consequently, joint analysis of gene expression and chromatin accessibility at the single-cell level can provide a more complete understanding of the gene-regulatory dynamics associated with human brain development.

To that end, we generated a transcriptomic and chromatin accessibility atlas, profiling 45,549 cells using multi-omic single-nucleus RNA-seq and ATAC-seq, across a broad developmental time frame that includes human fetal cortical plate, early postnatal, adolescent and adult specimens. We explored gene regulatory interactions by combining chromatin accessibility with gene expression within the same cells, and identified a subset of genes that are regulated by multiple nearby putative enhancers and have an important role in lineage determination during cortical development. To better understand the regulatory mechanisms driving neurogenesis, we performed pseudotime trajectory analysis and detected dynamic changes in chromatin accessibility preceding transcript production as a critical component of neuronal lineage commitment. We evaluated the enrichment of lineage-specific genes and chromatin accessible regions with genetic risk loci for neuropsychiatric disorders in order to explore their cellular ontogeny. Taken together, our data present a valuable resource for understanding the gene-regulatory dynamics associated with human brain development, and for prioritizing targets for further study as well as the generation of therapeutics to treat neurodevelopmental disorders.

## Results

### Single-nucleus gene expression and chromatin accessibility profiles revealed congruent cell types in the human cortex

We used the 10X Chromium Single Cell Multiome ATAC + Gene Expression kit to simultaneously profile the transcriptome (via snRNA-seq) and chromatin accessibility (via snATAC-seq) in twelve human neocortex samples from six developmental periods (early mid gestation fetal, late mid gestation fetal, infancy, childhood, adolescence and adulthood) (**Fig. 1A**; **table S1**). To confirm that the paired profiles were truly derived from the same cells, we first performed multi-omic profiling on two samples containing mixtures of human and mouse cell lines, and asked whether the co-assayed cells were consistently assigned to the same species labels. As expected, no doublets were identified and we observed that human and mouse reads were well separated based on the chromatin and transcriptome profiles of filtered cells (**fig. S1A**).

**Figure 1.**
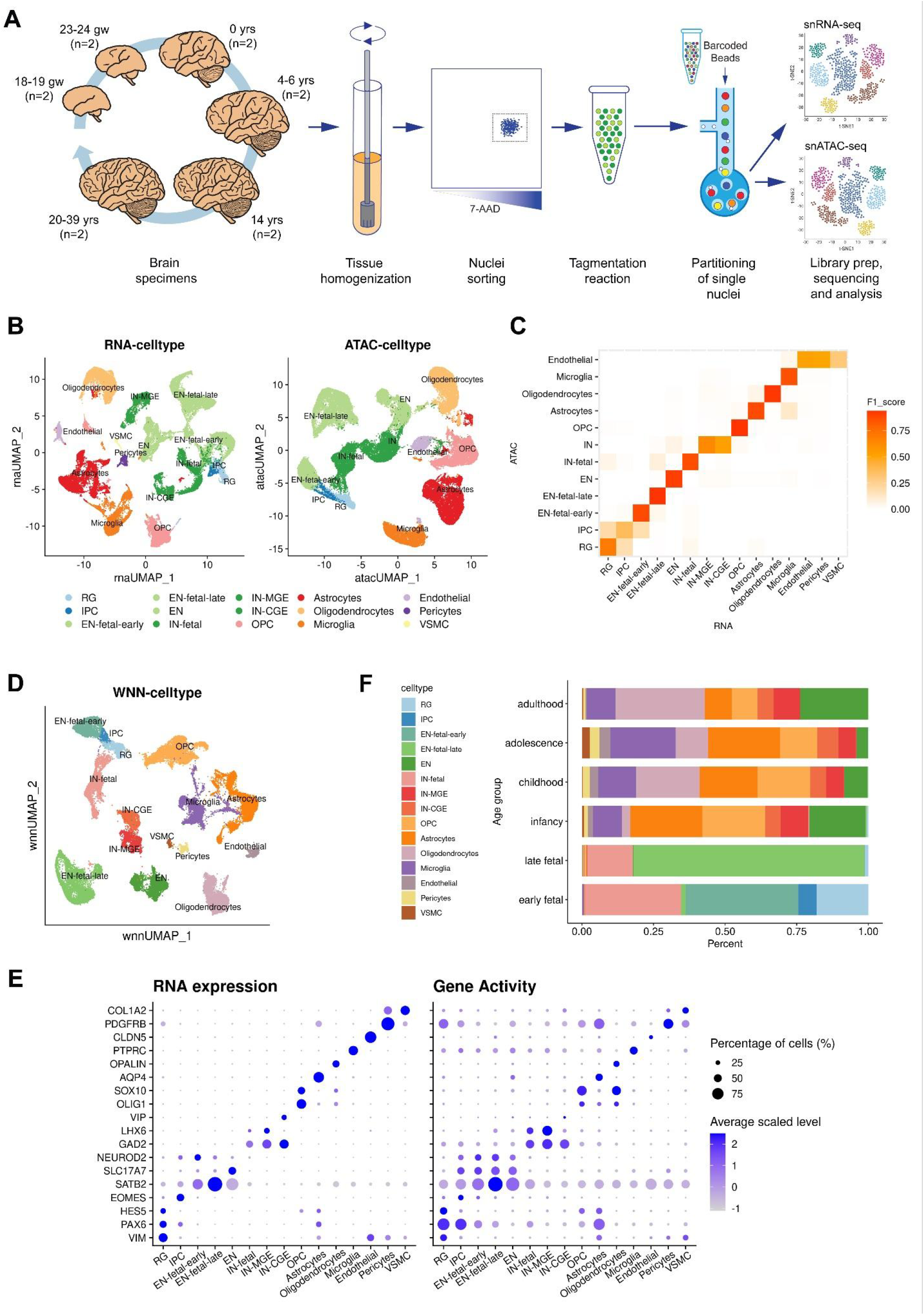
Joint single-cell profiling of RNA expression and chromatin accessibility of human neocortex. **(A)** Frozen human cortical brain specimens from 6 developmental time points were homogenized and purified by FANS prior to tagmentation and partitioning into single nuclei using the 10x Genomics platform. Libraries for snRNA-seq and snATAC-seq were prepared, sequenced and analyzed independently. **(B)** UMAP visualizations of single cells defined by RNA-seq and ATAC-seq data, respectively. Cell type annotations are derived from either modality independently. **(C)** Heatmap showing the concordance of cell memberships between the two clustering results, measured in F1 score. **(D)** UMAP visualization of single cells defined by integrating two modalities using WNN analysis. Cell type annotations are determined on the basis of marker genes. **(E)** Dot plot showing selected marker gene expression and chromatin-derived gene activity across cell types. **(F)** Proportions of cell types in each age group.

We then processed human neocortex samples, obtaining joint profiles of chromatin accessibility and gene expression from 45,549 out of 53,185 single nuclei that met quality control and filtering criteria (**Materials and Methods**). To assess the similarities and differences between the two modalities, we first clustered the RNA-seq and ATAC-seq data sets independently (**Fig. 1B**; **Materials and Methods**). Broadly, both modalities revealed the same major neocortical cell types and that cell identities assigned to RNA-seq and ATAC-seq derived cell types were highly congruent (**Fig. 1C**; adjusted Rand index [ARI] = 0.78).

Similar to previous multi-omic single-cell studies (*14, 15*), the independent modality analyses exhibit differences, primarily in the composition of cell populations in the fetal and postnatal stages (**fig. S1B**). On one hand, some cell types were broadly identified but not distinguishable in the ATAC-seq clustering results. For example, the medial ganglionic eminence (MGE)-derived and caudal ganglionic eminence (CGE)-derived inhibitory neuron subtypes were not distinguished; various stromal cell types with smaller population sizes, including endothelial cells, pericytes and vascular smooth muscle cells (VSMCs), were blended together. On the other hand, RNA-seq data showed insufficient power to identify progenitor cells, as evidenced by nearly 20% fewer detected radial glia (RG) and intermediate progenitor cells (IPCs) when compared with the ATAC-seq results (1,427 for RNA-seq vs. 1,743 for ATAC-seq), indicating that active gene-regulatory dynamics at different developmental stages might be better reflected in chromatin accessibility than in the transcriptome (*16*). These results motivated us to anticipate more comprehensive information about cell-type classifications by leveraging both modalities.

### Joint analysis of multi-omic data improved *de novo* taxonomy

We next performed joint clustering on the paired modalities of the same single cells using a weighted-nearest neighbor (WNN) analysis (*15*). WNN is an unsupervised method that generates an integrated representation of cellular identity by learning the information content of each modality. The WNN analysis results were in agreement with those derived from either single modality (ARI = 0.88 for RNA-seq, ARI = 0.86 for ATAC-seq), while the inferred relative modality weights varied across cell types (**fig. S1C**), reflecting the biological importance of each modality in determining cellular identity. The WNN analysis resulted in 28 clusters, including all the major and minor cell types in the human brain cortex, which were further grouped into 15 cell types (**Fig. 1D**; **Materials and Methods**). We confirmed that each cluster comprised cells from different samples (**fig. S1D**), suggesting that taxonomy was not determined by donor or other technical covariates.

Gene activity inferred by gene expression and chromatin accessibility of known cell type-specific markers consistently confirmed cluster identity (**Fig. 1E**; see lists of differentially expressed genes and accessible peaks in **table S2** and **table S3**; **Materials and Methods**). Specifically, we found neural progenitor cells expressing *PAX6*, including RG (1 cluster; *HES5, VIM*) and IPCs (1 cluster; *EOMES*). We also identified three subtypes of excitatory neurons (*SATB2, SLC17A7, NEUROD2*) representing different developmental stages, one enriched for cells from early fetal samples (‘EN-fetal-early’; 4 clusters), one for late fetal samples (‘EN-fetal-late’; 2 clusters), and the third for postnatal samples (‘EN’; 2 clusters). Similarly, there were three subtypes of inhibitory neurons identified (*GAD1, GAD2*), two of which represent MGE-derived (‘IN-MGE’; 1 cluster; *LHX6*) and CGE-derived (‘IN-CGE’; 1 cluster; *VIP, ADARB2*) subtypes in postnatal samples, while the remaining subtype was enriched in fetal samples (‘IN-fetal’; 1 cluster). The types of neurons that are distinct between fetal and postnatal human brain samples support previous findings (*17*). In addition, we observed clusters of major glial cell types in the neocortex, including oligodendrocyte progenitor cells (OPCs; 2 clusters; *OLIG1, SOX10*), astrocytes (3 clusters; *AQP4, GFAP*), oligodendrocytes (3 clusters; *MOBP, OPALIN*), microglia (4 clusters; *PTPRC, CX3CR1*), as well as endothelial cells (1 cluster; *CLDN5*), pericytes (1 cluster; *PDGFRB*) and VSMCs (1 cluster; *COL1A2*).

Sample-specific cell type composition varied significantly across developmental stages (**Fig. 1F**). In the four fetal samples, neuronal populations accounted for the vast majority of cells, whereas postnatal samples had much higher proportions of non-neuronal cells. The changing patterns of cell type composition were in line with the results from a previous deconvolution study using multiple bulk and single-cell datasets (*18*). Moreover, we found that most of the neural progenitors (91%), including the transient cell types of RG and IPCs, were only detected in the two early fetal samples (gestational week [GW] 18-19; **Fig. 1F, fig. S1E**), consistent with the fact that the bulk of neurogenesis in the human cerebral cortex has occurred by midgestation (at GW20) and these progenitor cells start disappearing or transforming with the completion of cortical development (*19, 20*). Notably, the results derived from joint analysis identified every cell type that was found in either single-omic analyses, while not losing power for detection of neural progenitors (1,736 by joint analysis vs. 1,743 by ATAC-seq alone vs. 1,427 by RNA-seq alone).

### *Cis*-regulatory associations between chromatin peaks and target genes

Multi-omic data offer the advantage to explore gene regulatory interactions by combining chromatin accessibility with gene expression within the same cells. Due to the sparsity of snATAC-seq and snRNA-seq data, we examined the relationships between the two modalities using pseudobulk aggregates rather than individual cells (*14, 16, 21*). We generated 500 pseudobulk samples by aggregating RNA-seq and ATAC-seq signals from similar cell types (**fig. S2A**; Methods). First, we sought to globally quantify the relative contribution of proximal (i.e., promoter) and distal (i.e., enhancer) chromatin accessibility to transcriptional variance. We applied a variance component model to the expression of each gene using the covariance of chromatin accessibility at promoter and enhancer regions as inputs, and corrected for donor and age effects by adding the inter-individual and inter-age-group covariance to the model (*22, 23*) (**Materials and Methods**). This approach does not model the relationship of each gene to its own promoters or enhancers, but instead models the genome-wide relationships to all enhancers or promoters. Our results suggested that more than 80% of expression variance was attributed to promoter and enhancer accessibility (**Fig. 2A**), indicating that transcriptional heterogeneity is broadly associated with the variation of chromatin accessibility. As control, we randomly permuted the dataset and, as expected, a minimal proportion of variance (< 1%) was explained by the epigenome in the shuffled analysis (**fig. S2B**). There was a small group of genes (n = 56) for which > 60% of the expression variance could be best explained by the inter-age-group covariance. Gene ontology (GO) enrichment analysis of these genes revealed enrichment in DNA-binding transcriptional activators (FDR q-value = 0.02; including known TFs like *SOX11, SOX4, NEUROD6, NR3C1*), suggesting the temporal role for these TFs in human brain development.

**Figure 2.**
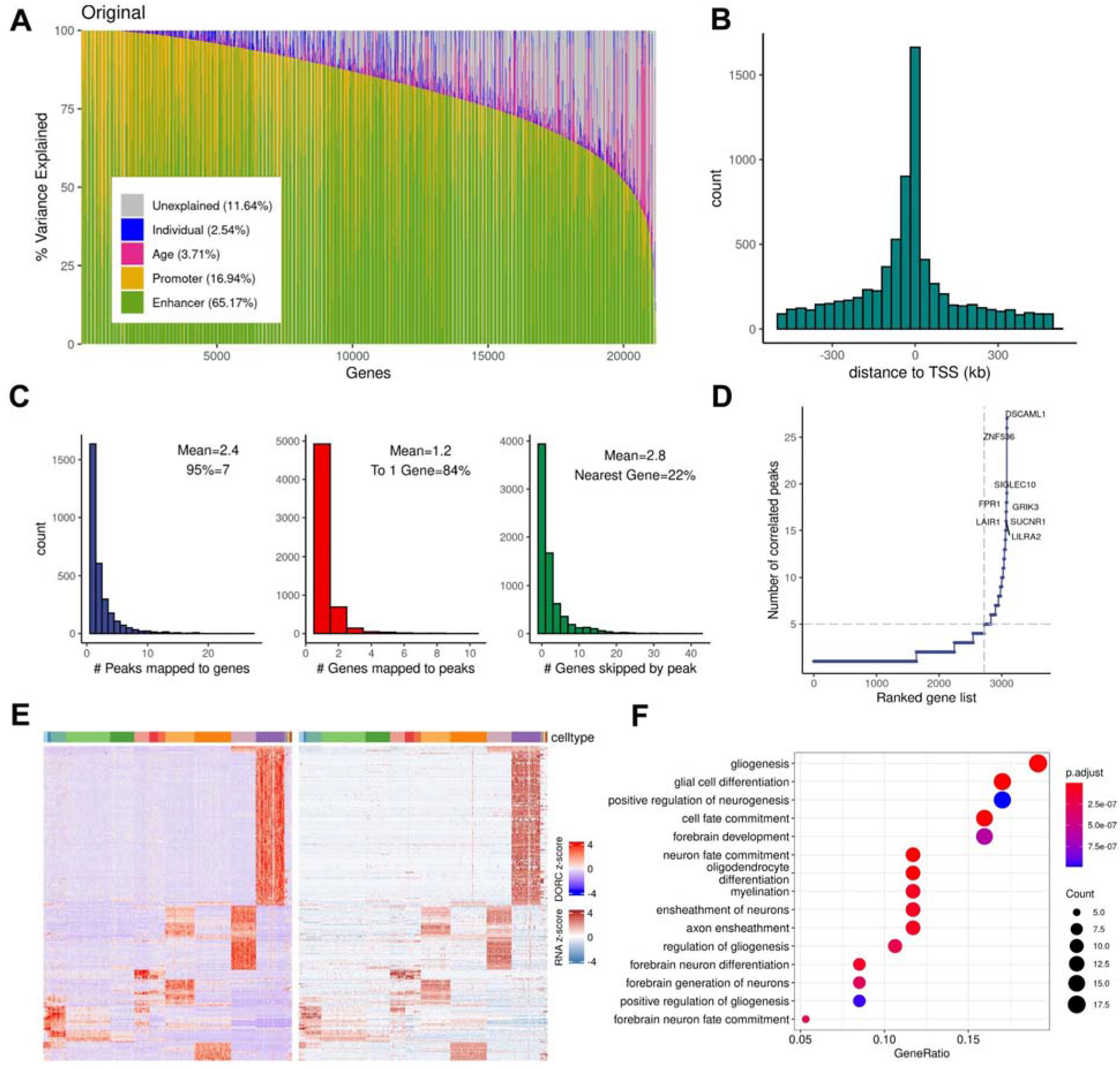
Global and local characterization of *cis* regulation patterns. **(A)** Variance component analysis showing chromatin accessibility explains variation in gene expression. Genes, in columns, are sorted by decreasing proportion of variance explained by the epigenome (enhancers and promoters), with the mean variance explained by each component shown in parenthesis. **(B)** Distribution of the distance from each peak to the transcription start site (TSS) of the linked gene. **(C)** Histograms showing (from left to right) distribution of the number of peaks significantly linked per gene; distribution of the number genes significantly linked per peak; distribution of the number of genes “skipped” by a peak to reach its linked gene. **(D)** The number of significantly linked peaks for each gene, with genes sorted in increasing order. **(E)** Heatmap showing chromatin accessibility and gene expression of the linked peak-gene pairs (rows, left aggregated peak accessibility, right linked gene expression) in the DORCs across 500 pseudobulk samples (columns, sorted in terms of cell types); values are z-score normalized. **(F)** Top 15 GO enrichment results for genes linked to DORCs.

With the aim of linking a regulatory element to its target gene(s), we next used a correlation-based approach to examine the pairwise relationships between chromatin accessibility and gene expression (**Materials and Methods**). This led to the identification of 7,291 significant peak-gene associations (within ±500kb around transcription start sites (TSSs), Spearman correlation coefficient |ρ| > 0.3, FDR-adjusted *P* value < 0.1; **table S4**), involving 3,082 unique genes. The majority (97.6%) of these links included peaks that were positively correlated with gene expression (**fig. S2C**). As expected, these associations were enriched in the vicinity of TSSs, and the correlations decayed exponentially with distance (**Fig. 2B**). Nevertheless, only 22% of the peak-gene links occur between an ATAC-seq peak and the nearest gene, indicating that the majority of predicted regulatory interactions skip at least one gene along the linear genome (**Fig. 2C**), demonstrating the shortcomings of purely applying the ‘nearest neighbor gene’ rule to define regulatory targets (*22, 24, 25*). The expression of most genes is, on average, correlated with at least two different peaks, while most peaks (84%) are predicted to interact with a single target gene (**Fig. 2C**). To validate the set of identified peak-gene links, we employed the ‘activity-by-contact’ (ABC) approach (*26*) (**Materials and Methods**) and compared them with the enhancer-promoter (E-P) interactions that were previously derived from the matched bulk brain tissues (*27*). We observed significantly higher ABC scores in the group of E-P interactions overlapping with the peak-gene links (*P* value < 2.2 × 10^−16^ by Wilcoxon test; **fig. S2D**), thereby providing further validation.

### Cell type specific *cis*-regulatory domains determine cell lineage during cortical development

To investigate the specificity of peak-gene associations across cell types and developmental stages, we assigned each interaction to the cell type with the highest average gene expression and chromatin accessibility. Peak-gene associations were strongest in the early developmental stage while they became diminished in more differentiated stages (**fig. S2E**). Specifically, RG-specific peak-gene links were the strongest across all cell types; in the group of neurons (either excitatory or inhibitory), which consist of samples from fetal to postnatal stages, we observed a clear weakening pattern of the associated links with developmental age. We defined a ‘pseudo-age’ for each cell type (**Materials and Methods**) and confirmed a significantly negative relationship with the median link strengths (Pearson’s *r* = -0.57, *P* value = 0.026; **fig. S2E**).

Despite the fact that most genes involved in peak-gene links were associated with one or two peaks, a subset of genes were associated with a relatively large number of peaks, suggesting orchestrated coregulation of the target gene activity by multiple factors that act upon a broad chromatin domain. In total, we identified 364 domains of regulatory chromatin (DORCs) (*14*) in which there are at least five significant peak-gene links associated with the same gene (**Fig. 2D**; **Methods**). In previous studies, it has been shown that DORCs are often associated with super-enhancers -- large clusters of enhancer regions that are known to play key regulatory roles in defining cell identity and are affected across multiple diseases (*28, 29*). Consistent with these studies, we found that DORCs identified here were also prominently overlapped with super-enhancers, which were identified by utilizing neuronal and glial ChIP-seq H3K27 acetylation data from human brain samples (*27*) (*P* value = 8.3 × 10^−68^ by hypergeometric test; **table S5**). For example, the DORC of the *DSCAML1* gene contained 27 peak-gene associations. The epigenetic dysfunction of this super-enhancer has been implicated in Alzheimer’s disease pathology (*30*).

Motivated by previous studies (*14, 31*), we hypothesized that DORCs are highly cell-type-specific. We defined a DORC score for each gene as the aggregated normalized counts from all peaks significantly associated with that gene (**Materials and Methods**). Covariation of chromatin accessibility and gene expression distinguished the identified cell types in both RNA-seq and ATAC-seq data (**Fig. 2E**), suggesting the cell-type specificity of DORC-gene links. Gene ontology (GO) analysis of the genes involved in the top decile of the peak-gene correlations in DORCs revealed strong enrichment of developmental processes in both neurons and glia (**Fig. 2F**), highlighting the important role of DORCs in cell fate determination during cortical development. Through comparison of neurons from different developmental stages (**table S6**), we found a higher number of DORCs specific to earlier stages (e.g., fetal versus postnatal, early fetal versus late fetal), suggesting a role in regulating early neurodevelopmental processes.

### Chromatin priming precedes gene expression during neuronal lineage commitment

Having identified various neuronal subtypes from early fetal cortical plate to adult cortical samples, we next utilized the paired multi-omic single nuclei profiles to infer the developmental dynamics of gene regulation throughout corticogenesis and neuron differentiation. We performed a pseudotime trajectory analysis by focusing on the neuronal populations (including RG, IPC, EN-fetal-early, EN-fetal-late, EN, IN-fetal, IN-MGE, and IN-CGE) and by anchoring the starting point in the RG cluster (**Materials and Methods**). Different cell types were properly laid on the inferred trajectories in terms of their developmental stages (**Fig. 3A, fig. S3A**), with the fetal-sample-specific neuronal populations located between the initial progenitor populations and the mature neurons from postnatal samples (i.e., EN, IN-MGE, IN-CGE). The developmental trajectories separated into EN lineage and IN lineages shortly after the starting point, and the IN lineage later split into IN-MGE and IN-CGE subtypes (**Fig. 3B**). The respective numbers of cells assigned to each of the three lineages are: EN lineage (14,146), IN-MGE lineage (5,728) and IN-CGE lineage (4,904).

**Figure 3.**
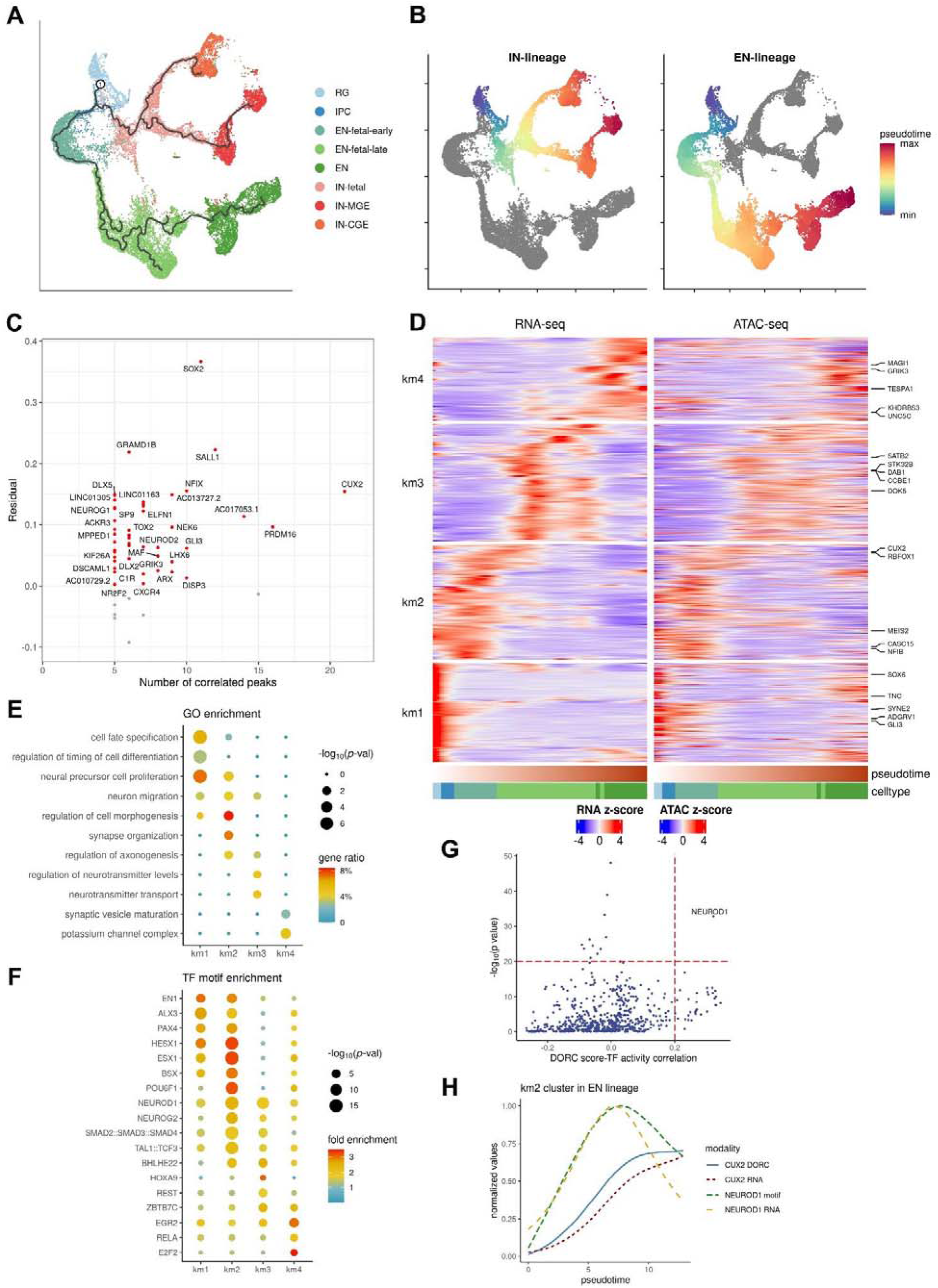
Trajectories of gene regulation during neuronal development. **(A)** Trajectories identified within the neuronal subpopulations, shown on the RNA gene expression coordinates (root node was annotated as ‘1’, cells were colored for annotated cell types). **(B)** Inferred pseudotime along the lineages for excitatory neurons (‘EN-lineage’) and inhibitory neurons (‘IN-lineage’), respectively. **(C)** Average residuals between chromatin accessibility and gene expression versus the number of significantly linked peaks for each gene involved in the DORCs identified within the neuronal populations. Positive and negative residuals are colored in red and gray, respectively. **(D)** Heatmap showing gene expression and DORC chromatin accessibility of the peak-gene links that significantly varied along the pseudotime for the EN lineage. Rows (genes) are clustered using k-means clustering (k=4), columns (cells) are ordered by pseudotime. The top five most differentially expressed genes in each cluster (‘km1/2/3/4’) are annotated. **(E)** Respective GO enrichment of genes represented in the four peak-gene link clusters of the EN lineage. **(F)** TF motifs enrichment of peaks represented in the peak-gene link clusters of the EN lineage. **(G)** *P* values of TF motif enrichment in km2 peaks plotted against Spearman correlation of TF motif activity with *CUX2* DORC score. **(H)** Lineage dynamics of *NEUROD1* motif activity and expression precede *CUX2* DORC chromatin accessibility and gene expression in the EN lineage, from the beginning to the end of the km2 stage, using the min-max normalized, smoothed values over pseudotime.

We repeated the peak-gene association analysis focusing on the neuronal populations (Methods), resulting in 1,638 significant associations involving 930 unique genes (**table S7**). Similarly, we defined 55 neuron-specific DORCs (associated with at least five peaks), which strongly overlapped with the DORCs that we defined using all cells (*P* value = 5.8 × 10^−51^, by hypergeometric test). GO analysis of the genes involved in these DORCs revealed a significant enrichment for neuron differentiation pathways as well as the overrepresentation of DNA-binding TFs, either activators or repressors (**table S8**).

Given the potentially tight regulation of DORC target genes by dynamic changes in chromatin accessibility during lineage commitment, we sought to explore whether chromatin accessibility at DORCs precedes gene expression. For each of the DORCs, we quantified a ‘residual’ by subtracting the corresponding gene expression value from the DORC score (*14*) (**Materials and Methods**). Remarkably, we observed that the residuals were typically positive (46 out of 55) across lineages (**Fig. 3C**), which reflected the lineage-priming of *cis*-regulatory elements, as the DORCs generally became accessible prior to onset of their associated gene’s expression. Furthermore, we found that the lineage-priming pattern became more robust for DORCs with a higher number of peaks, indicating higher confidence in the chromatin accessibility-primed states. Overall, these findings suggest that dynamic changes in chromatin accessibility is a critical component of neuronal lineage commitment, similar to previous observations during hair follicle differentiation (*14*).

### NEUROD1 induces CUX2 chromatin priming and gene expression during development of excitatory neurons

We next looked deeper into the peak-gene links on the EN lineage, which started from neuronal progenitors, including RGs and IPCs, and then differentiated into excitatory neuron subtypes specific to different developmental stages sequentially, from early fetal to late fetal and then to postnatal (**Fig. 3, A** and **B**). We found that the expression levels of over 87% of the linked genes (811/930) varied significantly along the pseudotime trajectory **(Materials and Methods**). We then grouped these genes into four clusters using k-means (km) clustering, each of which corresponding to a different developmental period (**Fig. 3D**). GO enrichment analysis on this gene set revealed the unique biological activities occurring during different time periods (**Fig. 3E**). Specifically, at the beginning of the trajectory (‘km1’), the linked genes were enriched in processes relating to cell fate specification, timing regulation of cell differentiation, and neural precursor cell proliferation. In the next early fetal period (‘km2’), the peak-gene interactions became associated with neuron migration, morphogenesis, synapse organization, and axonogenesis. Afterwards, in the late fetal (‘km3’) and postnatal stages (‘km4’), the excitatory neurons acquired the ability for neurotransmitter transport and regulation, indicating cell maturation.

The dynamic regulatory activities during the developmental transition of cell lineages are highly orchestrated by the spatiotemporal patterning of TFs. To identify TFs that control these dynamic regulatory activities, we performed TF motif enrichment analysis in the different clusters (**Materials and Methods**). TF motifs with an established function in cell differentiation and development were enriched in the earliest stage, including *EN1*, which has been implicated as a crucial mediator of dopaminergic subset specification (*32*), and *HESX1*, which has been identified as a hub gene for neural commitment (*33*) (**Fig. 3F**; **table S9**). In the intermediate stages (including early and late fetal), the associated peak-gene links were more enriched in motifs of neuronal TFs such as *NEUROD1, NEUROG2* and *BHLHE22*, suggesting the most active neurogenesis processes occur during these particular developmental periods. Fewer TF motifs were found enriched in the last postnatal peak-gene link cluster, including cell cycle regulators such as *E2F2*.

Cut Like Homeobox 2 (*CUX2*) was identified as a neuron-specific DORC gene, regulated by the highest number (n=21) of nearby putative enhancers (**Fig. 3C**), and as a marker for the second earliest stage in the EN lineage (‘km2’; **Fig. 3D**). This is consistent with the well-known function of *CUX2* as a neuron-specific TF regulating dendritic branching and synapse formation (*34*). Next, we investigated which TF(s) might activate *CUX2* enhancers by leveraging the correlation between the DORC score of *CUX2*, the TF motif activity (inferred from ATAC-seq) and the TF motif enrichment for the km2 cluster (**Materials and Methods**). The binding motif for the TF *NEUROD1* was strongly enriched in km2 chromatin accessible regions and *NEUROD1* activity was highly correlated with the *CUX2* DORC chromatin accessibility state (**Fig. 3G**). *NEUROD1* is essential for eliciting the neuronal development program and possesses the ability to reprogram other cell types into neurons (*35*). We next ordered single cells based on the inferred pseudotime for the km2 stage and identified a clear pattern where the activity of *NEUROD1* precedes the *CUX2* DORC chromatin state, followed by *CUX2* gene expression (**Fig. 3H**). Additionally, we found that, as *NEUROD1* activity decreases, the rate of *CUX2* expression slows down accordingly. These results suggest that *NEUROD1* is likely a key TF during early neurogenesis (*35*) to induce *CUX2* DORC accessibility followed by *CUX2* transcription.

### Repression of *NEUROD1* expression in cultured neural progenitor cells suppress *CUX2* expression

We sought to validate the predicted causal relationship between *NEUROD1* and *CUX2* by performing CRISPRi in cultured neural progenitor cells (NPCs) followed by RNAscope to directly image mRNAs in single cells. We transduced guide RNAs against both NEUROD1 and CUX2 into NPCs stably expressing dCas9-KRAB and then differentiated the NPCs using an established protocol (*36*). In the negative control experiment, in which cells were treated with a scrambled guide RNA, we observe that *CUX2* is widely expressed albeit at low numbers per cell, whereas *NEUROD1* expression is restricted to a smaller subset of cells but with a broader range of mRNA numbers per cell, including a fraction displaying strong bursts of transcription (**Fig. 4A**). We quantified the frequency distribution of fluorescent dots per nucleus for both genes at week 2 post-differentiation and found that *NEUROD1* expression is more variable than *CUX2* across the population, as measured by the fano factor ((variance/mean), CUX2 = 1.78, NEUROD1 = 17.02) (**Fig. 4B**).

**Figure 4.**
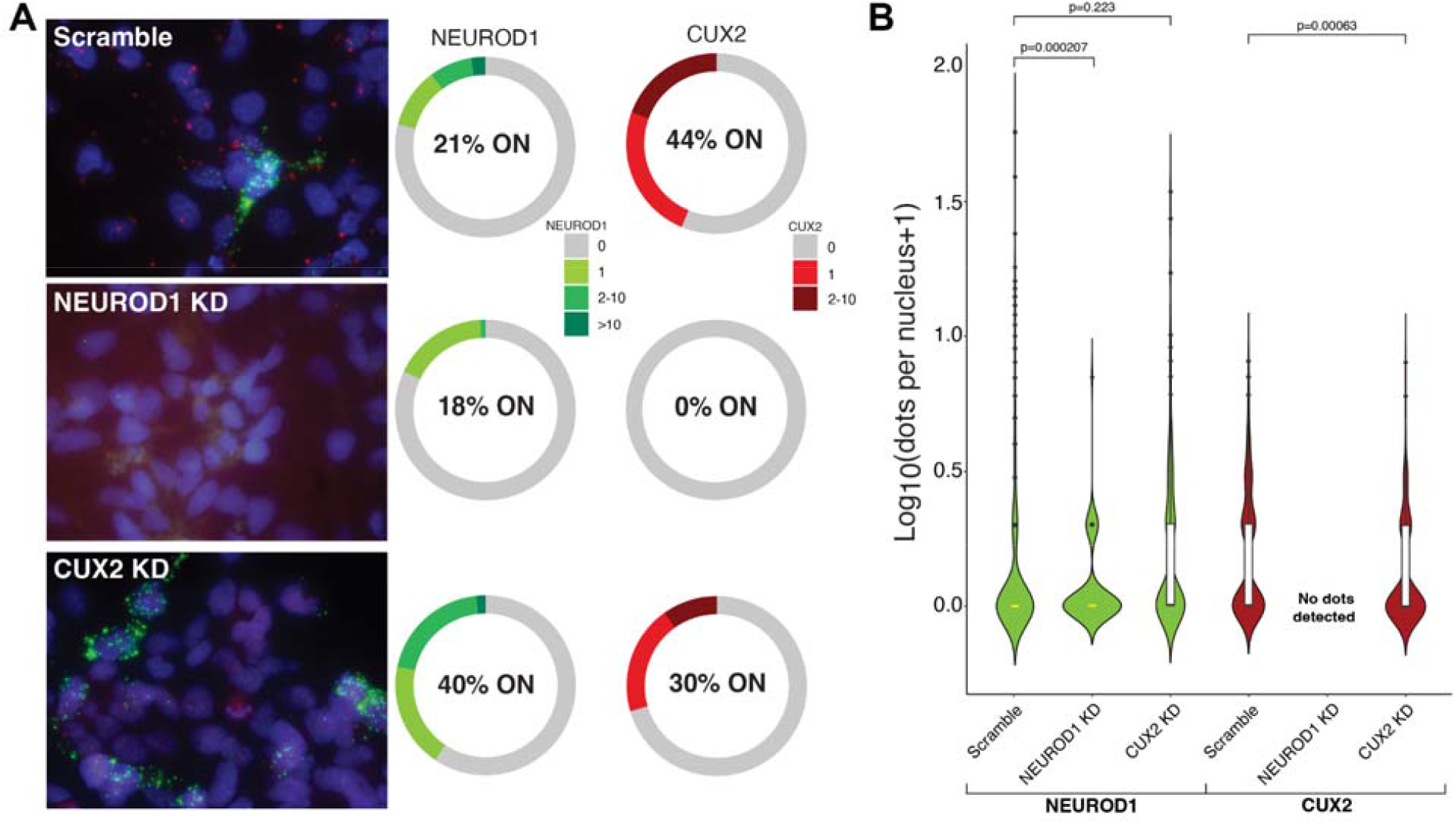
Assessment of the relationship between NEUROD1 and CUX2 in differentiating neural progenitor cells. **(A)** Maximum intensity projection images (from 200 nm z stacks obtained at 63x magnification) of CUX2 (red) and NEUROD1 (green) expression in NPCs 2 weeks post-differentiation treated with scrambled gRNA, a NEUROD1 specific gRNA and a CUX2 specific gRNA. Charts show frequency distributions of RNAscope dots per nucleus for CUX2 and NEUROD1 in cells treated with scrambled gRNA (n=444 cells), NEUROD1 specific gRNA (n=111 cells) and CUX2 specific gRNA (n=183 cells). % ON corresponds to % of nuclei with detectable RNAs. **(B)** Violin plots of nuclear RNA frequency distributions in all conditions. A negative binomial test was performed. The center line (yellow) indicates the median, the box shows the interquartile range, whiskers indicate the highest/lowest values within 1.5x the interquartile range.

Inactivation of NEUROD1 led to a down-regulation of *NEUROD1* mRNA compared to the control (*P* value = 0.0002 by negative binomial test), while CUX2 transcription is completely suppressed (**Fig. 4, A** and **B**). Given that around 80% of nuclei in the control NPCs do not show transcription of NEUROD1, the NEUROD1 promoter may be tightly repressed for long periods but allows for infrequent, strong bursts of transcription. In turn, inactivation of CUX2 with CRISPRi led to down-regulation of *CUX2* mRNA and a decrease in the proportion of cells expressing CUX2 compared to control (*P* value = 0.0006 by negative binomial test), without affecting NEUROD1 expression (*P* value = 0.223 by negative binomial test). Altogether, our data suggests that although NEUROD1 is expressed infrequently, it is required to maintain ongoing transcription of CUX2.

### Dissociation of risk loci for neuropsychiatric traits using single cell-derived marker genes and peaks

Despite the notable progress in exploring the genetic causes of neuropsychiatric disorders, their underlying molecular mechanisms are still not fully understood (*37*). To reveal whether disorder-associated variants are enriched in a particular cell type or developmental stage, we used linkage disequilibrium-aware approaches (**Materials and Methods**) (*38, 39*) to assess the overlap between a collection of relevant GWAS studies and lineage-defining genes and chromatin peaks derived from our multi-omic single cell data. By analyzing 9 neuropsychiatric and 3 unrelated control traits in 15 cell types, we identified 33 and 28 significant associations in cell type-specific chromatin accessibility and transcriptome data, respectively (**Fig. 5A, table S10** and **S11**). We observed a high overlap of significant cell-type - GWAS trait pairs (20/41 of significant pairs are shared; Spearman correlation of all pairs ρ=0.62), suggesting that both modalities report reliable and informative associations.

**Figure 5.**
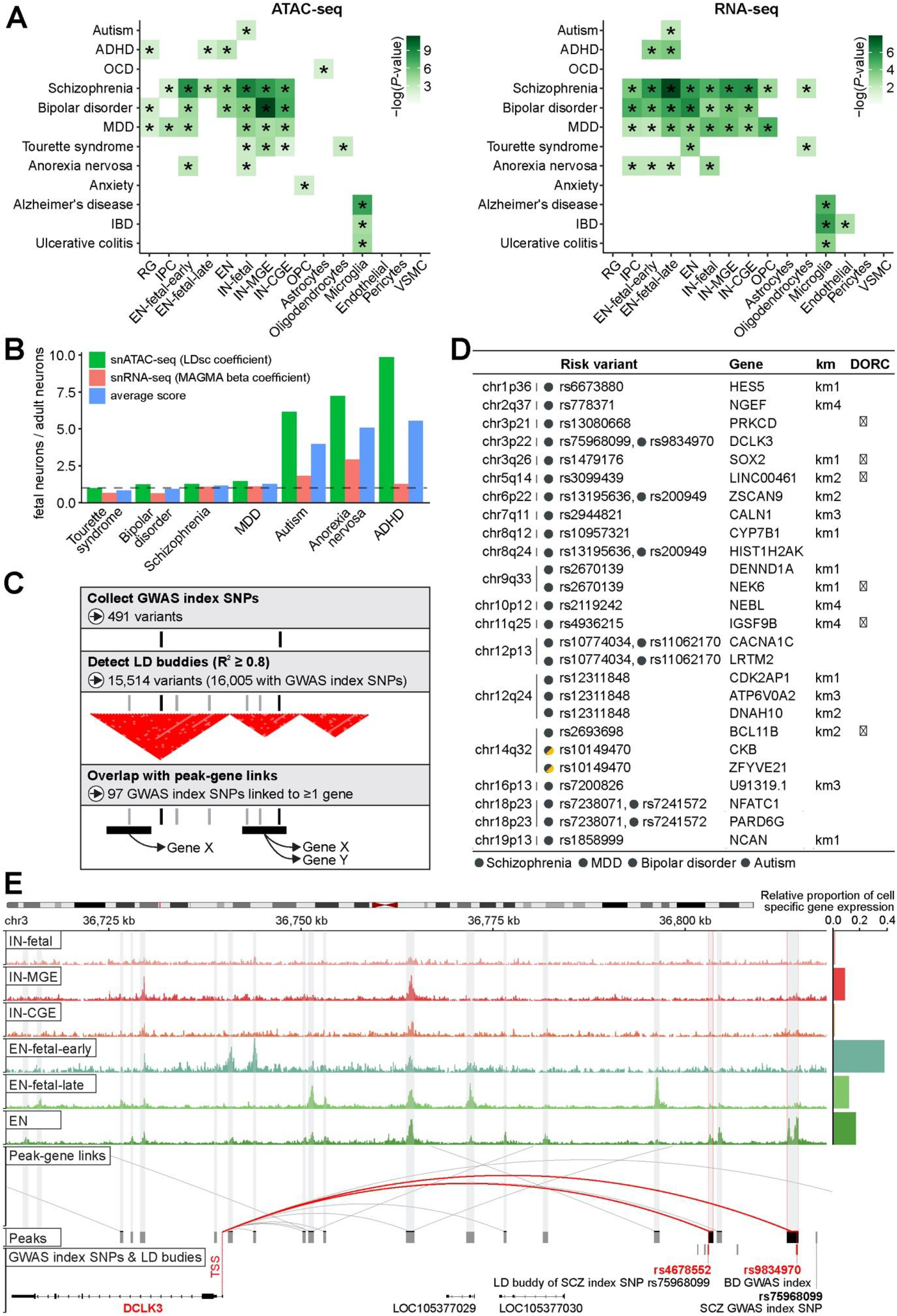
Mapping of risk variants associated with neuropsychiatric traits to causal genes using single cell-derived marker genes and peaks. **(A)** Heritability enrichment of brain cell types in neuropsychiatric disorders and unrelated control traits. Heatmaps highlight significant co-localization of GWAS-derived common genetic variants with cell-specific open chromatin regions in snATAC-seq data (left panel) and cell marker genes in snRNA-seq data (right panel) (Methods). “*”: significant after correction across all tests (FDR<0.05). **(B)** Comparison between fetal and adult neuronal signals in selected neuropsychiatric disorders (traits need to be enriched in either fetal or adult category; therefore, OCD and Anxiety were not involved). Fetal and adult neurons are represented by peaksets / genesets compiled from unions of the top 2,500 / 500 the most cell-specific peaks / genes from each fetal neuron (i.e., EN-fetal-early, EN-fetal-late, and IN-fetal) and adult neuron (i.e., EN, IN-CGE, and IN-MGE) category. To calculate the ratio “fetal neurons / adult neurons” (y-axis), we used LDsc regression coefficient (snATAC-seq) and MAGMA beta coefficients (snRNA-seq); joint score is an average of snATAC-seq and snRNA-seq scores. **(C)** Schematic of the overall strategy to connect risk variants associated with neuropsychiatric disorders to their causal genes. (**Methods**). **(D)** Subset of candidate causal genes for risk variants that are either prioritized in two disorders, or show significantly altered expression along the developmental trajectory of the neuronal lineage (km1/2/3/4; full list of causal genes in **Table S12**). **(E)** Normalized snATAC-seq-derived pseudobulk tracks demonstrating the complex cell-specific regulation of the DCLK3 gene that is predicted to be the causal gene for SCZ and BD GWAS risk variants (rs75968099 and rs75968099).

Consistent with previous studies, schizophrenia (SCZ), bipolar disorder (BD), attention deficit hyperactivity disorder (ADHD) and major depressive disorder (MDD) are enriched in multiple neuronal subtypes (*40–42*). However, to our knowledge, our results identified several associations that have not yet been described by genetic data (see Discussion), including oligodendrocytes for Tourette syndrome (TS), astrocytes for obsessive-compulsive disorder (OCD), OPCs for anxiety and fetal neurons for anorexia nervosa (AN). For non-psychiatric, immune-related traits, including Alzheimer’s disease (AD), ulcerative colitis, and inflammatory bowel disease (IBD) we only observed enrichment for microglia, thus further strengthening the credibility of our results. To dissect the temporal specificity in neuropsychiatric diseases, we compared fetal and adult neuronal enrichment using both epigenome and transcriptome data. We found a high concordance between enrichment of both assays, allowing us to classify ADHD, AN and autism as being more strongly associated with fetal neuronal stages, while, for TS, BD, SCZ and MDD, we found an equal contribution of both fetal and adult neuronal stages (**Fig. 5B**).

We next aimed to nominate the candidate functional genes for disease-associated loci (**Fig. 5C, Materials and Methods**). First, we collected a set of 491 genome-wide significant variants associated with neuropsychiatric traits (P<5×10^−8^) and extended it to 16,005 variants based on the presence of high linkage disequilibrium (LD; R^2^≥0.8). We overlapped putative disease-relevant variants (index SNP and LD buddy) with the peaks that demonstrate significant peak-gene associations to pinpoint at least one gene under regulation for 97 genome-wide significant loci (**table S12**). Out of 152 genes mapped to those 97 loci, 7 were linked to two disease traits simultaneously and 17 genes were shown to have significantly altered expression along the pseudotime trajectory of neuronal lineage specification (km1/2/3/4) (**Fig. 5D**). While the original GWAS studies usually nominate several plausible gene targets for each disease-relevant locus, their prioritization is mostly based on imprecise distance-based annotation. Using our approach, we were able to refine their predictions and, in some cases, nominate novel genes. One example of a replicated finding is the association of *DCLK3* (encoding a neuroprotective kinase) with both SCZ and BD, which was previously observed by TWAS (*43*) and eQTL approaches (*44*) (**Fig. 5E**). Notably, this association is derived from the overlap of putative disease-relevant variants with two distinct peaks, both predominantly accessible in adult excitatory neurons.

## Discussion

We generated multi-modal chromatin accessibility and gene expression data in the human cortex across 6 broad developmental time-points from fetus to adult. Joint analysis of 45,549 individual nuclei facilitated the identification of genes and cis-regulatory elements with fundamental roles in lineage determination. By using the covariance of chromatin accessibility at promoter and enhancer regions as inputs, we show that the majority of expression variance was attributed to promoter and enhancer accessibility, indicating that gene expression is broadly associated with chromatin accessibility. Moreover, through comparison of neurons from different developmental stages, we found that there were more DORCs specific to earlier stages (e.g., fetal versus postnatal, early fetal versus late fetal), suggesting a role for chromatin reorganization in regulating early neurodevelopmental processes. The temporal nature of our data allowed us to examine neural trajectories across 4 broad developmental phases. The first of these contains genes involved in cell fate specification, differentiation, and NPC proliferation. The second cluster specifies genes involved in neuron migration, morphogenesis, synapse organization, and axonogenesis, while the 3rd and 4th clusters contain genes associated with neurotransmitter transport and regulation. As an example, we chose to focus on CUX2, a TF involved in synaptogenesis that is expressed in the 2nd cluster. CUX2 expression coincides with a number of nearby open chromatin regions containing putative enhancers, among which are binding sites for NEUROD1, a well known pioneer factor involved in neuronal cell fate specification. It has recently been shown that overexpressing NEUROD1 in astrocytes can convert them into neurons, suppressing the astroglial gene expression program while upregulating neuronal genes, including CUX2 (*45*). Thus, we hypothesized that NEUROD1 might activate CUX2 during early neural development, and subsequently showed that inactivation of NEUROD1 in cultured NPCs led to a complete suppression of CUX2, while inactivation of CUX2 did not affect expression of NEUROD1.

Lineage specific genes and chromatin accessible regions are enriched for risk loci associated with neuropsychiatric traits, and implicate 152 putative risk genes in a range of disorders, including SCZ, BD, ADHD and MDD. SCZ, BD, ADHD, and MDD are enriched in multiple neuronal subtypes, consistent with previous studies (*40–42*). Beyond already known associations between various cell types and disease, our results identified several associations that, to our knowledge, have not been described previously. First, TS was found to be enriched in oligodendrocyte cells in both epigenome and transcriptome assays. The critical role of oligodendrocytes is supported by tract-based spatial statistics measurements of TS patients, indicating a reduced fractional anisotropy that reflects deficits in axonal myelination (*46*). Second, OCD was enriched in astrocytes. While the literature supporting this relationship is more established for the striatum (*47*), the involvement of the prefrontal cortex was previously studied through an astrocyte-specific deletion of glutamate transporter 1 (*48*) and vesicular monoamine transporter 2 (*49*), both resulting in OCD-like behavior. Third, anxiety was enriched in OPCs, further implicating the well-established role for aberrant myelination in neuropsychiatric disorders (*50*). This relationship was emphasized by a recent study linking anxiety-like behavior in a mouse model of cuprizone-induced demyelination which displays impaired OPC differentiation (*51*). Lastly, we report the enrichment of fetal neurons in AN. While this disorder phenotypically manifests in adolescence or early adulthood, a number of studies suggest substantial changes during earlier stages of development (*52, 53*). Furthermore, significant differences in gene expression were previously measured between AN case and control subjects using hiPSC-derived cortical neurons (*54*) that are known to resemble fetal, rather than adult, brain cells (*55*).

In conclusion, we generated an atlas of gene expression and chromatin accessibility in single nuclei from 6 developmental time-points that provides additional insights on cell fate determination in the human cerebral cortex and on the molecular basis of neuropsychiatric disease. We present our data as an interactive web browser that can be utilized by the scientific community to explore spatiotemporal alterations in gene expression in development and disease.

## Materials and Methods Summary

Nuclei were isolated from frozen human cortical brain specimens from 6 developmental time-points (fetal 18-20 gestational weeks (GW) (n=2), 23-24 GW (n=2), 0 years (n=2), 4-6 years (n=2), 14 years (n=2) and 20-39 years (n=2)) and subjected to fluorescence-activated nuclear sorting (FANS). Purified nuclei were processed using the Chromium Next GEM Single Cell Multiome ATAC-seq and Gene Expression protocol (10x Genomics). Resulting libraries were sequenced using the Novaseq platform (Illumina), obtaining 100 bp paired-end reads that were aligned by cellranger-arc (v.1.0.0). MACS2 (*56*) was used to call peaks. We used Seurat v4.0 (*15*) to construct a weighted nearest neighbor (WNN) graph and shared nearest neighbor graphs as well as to find differentially expressed genes for each identified call type. Differentially accessible peaks were identified by Signac v1.1.0 (*57*). We applied the variance component analysis (*22, 23*) to quantify the proportion of gene expression variation that is attributable to promoter, enhancer and individual covariance. We used chromVAR (*58*) to perform transcription factor analysis for all DNA motifs from the JASPAR 2020 database (*59*). For each cell type, we created psedobulk populations and used them to calculate correlations between peaks and nearby genes (within 0.5Mbp window), thus reconstructing gene regulatory associations. Pseudobulk populations were also used to define ‘pseudo-age’ for each cell type based on the proportion of cells found in the six different developmental stages. Monocle3 (*60*) was used to construct the trajectories across neuronal populations and to find differentially expressed genes on the trajectory of a specific lineage. We applied LD-score score partitioned heritability (*39*) (snATAC-seq) and MAGMA (*38*) (snRNA-seq) to investigate whether the cell specific peaks and genes might play a role in disease, and quantified their co-localization with common risk variants from 53 GWAS studies. We examined the relationship between NEUROD1 and CUX2 via CRISPR mediated knock down in differentiating NPCs, followed by RNAscope.

## Acknowledgments

We thank the patients and families who donated material for these studies. Brain tissue for the study was obtained through the NIH Neurobiobank from the following brain bank collections: The Mount Sinai/JJ Peters VA Medical Center NIH Brain and Tissue Repository and the University of Maryland Brain and Tissue Bank. We thank the computational resources and staff expertise provided by the Scientific Computing at the Icahn School of Medicine at Mount Sinai. We thank Dr. Pengfei Dong, Dr. Xinyi Wang and other members in the Roussos Lab for helpful discussion. We thank the computational resources and staff expertise provided by the Scientific Computing of the Icahn School of Medicine at Mount Sinai.

## Author contributions

P.R. conceived of and designed the project. J.F.F. and P.R. designed experimental strategies for single-nucleus profiling of human postmortem tissue. N.M.T. performed tissue dissections for fetal specimens. J.F.F. generated single-nucleus multi-ome data. S.R., J.M.V., O.L., A.L., K.J.B. and J.F.F. performed validation experiments. K.Z., J.B., T.C., D.L., G.C.Y. and P.R. designed analytical strategies. K.Z. conducted initial bioinformatics, sample processing and quality control for single-nucleus data. K.Z. developed the computational scheme and performed the downstream analysis. J.B., T.C and C.C. performed the GWAS enrichment analysis and mapping of GWAS loci to causal genes. Data analysis was supervised by D.L., G.C.Y. and P.R. K.Z., J.B., S.R., G.C.Y., J.F.F and P.R. wrote the manuscript with input from all authors.

## Data and materials availability

Raw data (FASTQ files) and processed data (BigWig files; coordinates of open chromatin regions; count matrices), were deposited in NIH GEO under accession no. GSE204684. UCSC genome browser tracks for cell type pseudobulks are available at https://labs.icahn.mssm.edu/roussos-lab/3dg_dual_assay/. Single cell data can be further inspected at the Broad Institute Single Cell Portal under accession no SCP1859. Code used throughout this study is available upon reasonable request from the corresponding authors.

## Supplementary Materials

Materials and Methods

Figs. S1 to S3

Data S1 to S12

