## Supplementary Material for "Multi-omic profiling of the developing human cerebral cortex at the single cell level"

**This PDF file includes:**

Materials and Methods

Figs. S1 to S3

Captions for Data S1 to S12

**Other Supplementary Materials for this manuscript include the following:**

Data S1 to S12

### Materials and Methods

### Materials

| **Reagent** | **Vendor** | **Cat#** |
| --- | --- | --- |
| Accutase | Innovative Cell Technologies | AT104 |
| DMEM (Dulbecco’s Modified Eagle Medium) | Gibco | 11965092 |
| RNAse Free PBS | Fisher Scientific | BP24384 |
| Corning Cell Culture Buffer: Dulbecoo's Phosphate-Buffered Salt Solution 1x | Fisher Scientific | MT21030CV |
| Recombinant RNAse Inhibitor | Takara | 2313A |
| BSA | Miltenyi Biotec | 130-091-376 |
| Tween-20 | Sigma-Aldrich | P1379 |
| Countess cell counting chamber slides | Invitrogen | C10283 |
| Trypan Blue | Gibco | 15250-061 |
| QuBit RNA HS Assay Kit | Fisher Scientific | Q32855 |
| Cell strainer | Sigma-Aldrich | BAH13680040 |
| QIAzol Lysis Reagent | Qiagen | 79306 |
| Chloroform | Sigma-Aldrich | C2432 |
| lentiGuide-Hygro-mTagBFP2 | Addgene | Cat# 99374 |
| 10X T4 DNA Ligase buffer | New England Biolabs | B0202S |
| T4 PNK | New England Biolabs | M0201L |
| Quick Ligation Buffer (2X) | New England Biolabs | M2200L |
| BsmB1 v2 | New England Biolabs | R0739L |
| T7 DNA Ligase | New England Biolabs | M0318S |
| High Efficiency NEB 10-beta Competent *E. coli*. | New England Biolabs | C3019H |
| QIAprep Spin Miniprep kit | Qiagen | 27106 |
| Formaldehyde solution (37 wt.% in H_2_O, contains 10-15% Methanol as stabilizer) | Sigma-Aldrich | 252549 |
| RNAscope Probe- Hs-NEUROD1-C2 | Advanced Cell Diagnostics | 437281-C2 |
| RNAscope Probe- Hs-CUX2-C3 | Advanced Cell Diagnostics | 425581-C3 |
| RNAscope H_2_O_2_ and Protease Reagents | Advanced Cell Diagnostics | 322381 |
| RNAscope Multiplex Fluorescent Detection Reagents V2 | Advanced Cell Diagnostics | 323110 |
| RNAscope Wash Buffer Reagents | Advanced Cell Diagnostics | 310091 |
| ImmEdge™ Hydrophobic Barrier Pen | Advanced Cell Diagnostics | 310018 |
| RNAscope Multiplex TSA Buffer | Advanced Cell Diagnostics | 322809 |
| RNAscope Probe Diluent | Advanced Cell Diagnostics | 300041 |
| Opal 570 Reagent Pack | Akoya Biosciences | FP1488001KT |
| Opal 690 Reagent Pack | Akoya Biosciences | FP1497001KT |
| Nunc™ Lab-Tek™ II CC2™ Chamber Slide System (8-well) | Thermo Fisher | 154941 |
| 24 x 50 mm microscope cover glass (thickness No.1) | Fisher Scientific | 12-545F |
| Prolong Gold Antifade Reagent | Thermo Fisher | P36934 |

**Methods**

#### Description of post-mortem brain samples

Fetal brain samples were collected from de-identified prenatal autopsy specimens without neuropathological abnormalities at the Icahn School of Medicine at Mount Sinai (MSSM). The cortical plate was dissected fresh from the anterior frontal lobe of anatomically intact brain specimens. Young and old brain samples were accessed through the NIH NeuroBioBanks at the University of Miami Brain Endowment Bank and Mount Sinai Brain Bank, respectively. All neuropsychological, diagnostic, and autopsy protocols were approved by the respective Institutional Review Boards.

#### Isolation and fluorescence-activated nuclear sorting (FANS)

All buffers were supplemented with RNAse inhibitors (Takara). Four samples were processed in parallel. 25mg of frozen postmortem human brain tissue was homogenized in cold lysis buffer (0.32M Sucrose, 5 mM CaCl_2_, 3 mM Magnesium acetate, 0.1 mM, EDTA, 10mM Tris-HCl, pH8, 1 mM DTT, 0.1% Triton X-100) and filtered through a 40 µm cell strainer. The flow-through was underlaid with sucrose solution (1.8 M Sucrose, 3 mM Magnesium acetate, 1 mM DTT, 10 mM Tris-HCl, pH8) and centrifuged at 107,000 g for 1 hour at 4°C. Pellets were resuspended in PBS supplemented with 0.5% bovine serum albumin (BSA).

Prior to FANS, volumes were brought up to 250 µl with PBS and 7AAD (Invitrogen) was added according to manufacturer’s instructions. 7AAD positive nuclei were sorted into tubes pre-coated with 5% BSA using a FACSAria flow cytometer (BD Biosciences).

#### Multiome ATAC-seq and Gene Expression library preparation

Following FANS, nuclei were subjected to 2 washes in 200 µl wash buffer (10x Genomics), after which they were re-suspended in 30 µl nuclei buffer (10x Genomics) and quantified (Countess II, Life Technologies). 8000 nuclei from each sample were subjected to the Chromium Next GEM Single Cell Multiome ATAC-seq and Gene Expression protocol (10x Genomics), according to manufacturer’s instructions. Resulting libraries were quantified using the KAPA library quantification kit (KAPA Biosystems) and fragment sizes determined by Tapestation (Agilent). All libraries were sequenced at New York Genome Center (NYGC) using the Novaseq platform (Illumina), obtaining 100 bp paired-end reads.

#### Quantification of Multiome ATAC-seq and Gene Expression

Fastq alignment, filtering, barcode counting, peak calling and counting of both ATAC-seq and gene expression molecules were performed with cellranger-arc (v.1.0.0).

#### Quality control and data processing

We processed the outputs of Cell Ranger ARC using Seurat v4.0 [(*15*)](https://sciwheel.com/work/citation?ids=11129215&pre=&suf=&sa=0) and Signac v1.1.0 [(*57*)](https://sciwheel.com/work/citation?ids=11949939&pre=&suf=&sa=0) to create a multi-omic Seurat object with paired gene expression and DNA accessibility profiles for each sample. For chromatin accessibility, we used MACS2 [(*56*)](https://sciwheel.com/work/citation?ids=57981&pre=&suf=&sa=0) as implemented in the function CallPeaks in Signac to call peaks from the fragment files. We removed any peaks on nonstandard chromosomes overlapping annotated genomic blacklist regions from the hg38 genome [(*61*)](https://sciwheel.com/work/citation?ids=7132831&pre=&suf=&sa=0). We then quantified fragment counts for each peak, per cell, using the FeatureMatrix function in Signac. Per-cell quality control metrics for each modality were computed, including mitochondrial percentage, nucleosome signal and transcriptional start site (TSS) enrichment score. Next, we combined the individual Seurat objects for all the samples into one single object using the merge function in Seurat. We then performed quality control based on metrics for both modalities by retaining cells with total RNA-seq count > 200 and < 50,000, total ATAC-seq count > 200 and < 100,000, mitochondrial percentage < 5%, nucleosome signal < 3 and TSS enrichment score > 1. Genes or peaks which are detected in < 10 cells are removed. As a result, 45,549 out of 53,185 single cells are retained, with 5,887 RNA UMI counts and 18,066 unique ATAC-seq fragments (29.3% fragments overlapping peaks) on average.

We normalized the gene expression count data using SCTransform [(*62*)](https://sciwheel.com/work/citation?ids=8000403&pre=&suf=&sa=0) and then performed principal component analysis (PCA) using the RunPCA function in Seurat. The first 30 principal components (PCs) were used in the downstream analysis. For the chromatin accessibility data, we performed latent semantic indexing (LSI) for dimension reduction [(*63*)](https://sciwheel.com/work/citation?ids=5611447&pre=&suf=&sa=0). We first normalized the ATAC-seq peaks using the log-TF version of the term frequency-inverse document frequency (TF-IDF) transformation by using the RunTFIDF function in Signac (parameter setting: ‘method = 3’). The top 25% most frequently observed peaks of the TF-IDF matrix were then selected by the FindTopFeatures function for singular value decomposition (SVD) using the RunSVD function in Signac. The top ten LSI components, excluding the first, were used in downstream analysis. The first LSI component was excluded because it typically captures sequencing depth (technical variation), as evidenced by the high correlation with the total number of counts for the cells (> 0.5). We also created a gene activity matrix inferred from ATAC-seq by using the GeneActivity function in Signac, which assesses chromatin accessibility at gene body and promoter regions.

#### Clustering and visualization

We performed graph-based clustering using the reduced dimensions which we selected for both assays, as described above. For joint multi-omic analysis, we used the function FindMultiModalNeighbors in Seurat v4.0 to construct a weighted nearest neighbor (WNN) graph by taking as input two dimensional reductions computed for each modality [(*15*)](https://sciwheel.com/work/citation?ids=11129215&pre=&suf=&sa=0). WNN identifies the nearest neighbors for each cell based on a weighted combination of two modalities. For single modality assay, we used the function FindNeighbor to construct a shared nearest neighbor graph. We then applied the Smart Local Moving algorithm [(*64*)](https://sciwheel.com/work/citation?ids=4951223&pre=&suf=&sa=0) on the derived graphs by using the function FindClusters (parameter settings: ‘algorithm = 3’ and ‘resolution = 0.2’).

For 2-dimensional visualization, we performed uniform manifold approximation and projection (UMAP) [(*65*)](https://sciwheel.com/work/citation?ids=6094688&pre=&suf=&sa=0) as implemented in the function RunUMAP in Seurat on the PCs and LSI components for gene expression and DNA accessibility data, respectively. For joint data visualization, UMAP was computed using the multi-omic WNN graph (parameter setting: nn.name = ‘weighted.nn’).

To find markers (genes or peaks) for each identified cell type, we used the function FindAllMarkers (parameter settings: ‘only.pos = TRUE’). For differentially expressed genes, we used additional parameters (‘min.pct = 0.2’, ‘min.diff.pct = 0.1’) to ensure the markers are sufficiently expressed in the corresponding cell group. For differentially accessible chromatin regions, we used a logistic regression model and added the total number of fragments as a latent variable, as suggested in the Signac tutorial (by specifying additional parameters: min.pct = 0.05, test.use = “LR”, latent.vars = “atac_peak_region_fragments”). Genes or peaks with adjusted *P* value (using bonferroni correction) < 0.05 were retained as the cell-type-specific markers.

#### Variance component modeling of gene expression

We performed variance component analysis to characterize how much gene expression variability could be explained by the patterns of chromatin covariance. We selected genes and peaks which are detected in at least 10% of total pseudobulk samples. Then, for the expression vector of each gene *g*, we fit the following variance component model as suggested in previous studies [(*22*, *23*)](https://sciwheel.com/work/citation?ids=13683793,6327580&pre=&pre=&suf=&suf=&sa=0,0):

*Y_g_* ~ $N\left( 0,P\sigma_{p}^{2}+E\sigma_{e}^{2}+I\sigma_{i}^{2}+A\sigma_{a}^{2}+U\sigma_{u}^{2} \right)$

where *Y_g_* is the centered and scaled normalized gene expression vector of gene *g*, *P* and *E* are sample-sample correlation matrices computed by chromatin accessibility in promoter (defined as within -1000 to +100 bp from TSSs) and enhancer regions (defined as within ±500 kb around TSSs), respectively; *I*, *A* and *U* capture the per-individual, per-age-group and noise terms, respectively. The values of the variance parameters of the model were estimated by the average information restricted likelihood estimation (AIREML; ‘gaston’ R package). To determine the proportion of the variance explained by each variance component, we generated a vector *V_g_*  which, by definition, sums to 1:

*V_g_* =$\langle\frac{\sigma_{p}^{2}}{\sigma^{2}},\frac{\sigma_{e}^{2}}{\sigma^{2}},\frac{\sigma_{i}^{2}}{\sigma^{2}},\frac{\sigma_{a}^{2}}{\sigma^{2}},\frac{\sigma_{u}^{2}}{\sigma^{2}}\rangle$, where $\sigma^{2}=\sigma_{p}^{2}+\sigma_{e}^{2}+\sigma_{i}^{2}+\sigma_{a}^{2}+\sigma_{u}^{2}$

For example, the proportion of the variance in expression for gene *g* explained by the promoters would be represented by the first element in the *V_g_* vector.

#### Transcription factor motif enrichment analysis

We tested a set of peaks of interest for overrepresentation of each DNA motif from the JASPAR 2020 database [(*59*)](https://sciwheel.com/work/citation?ids=7853654&pre=&suf=&sa=0) by using the FindMotifs function in Signac. The resulting *P* values are FDR adjusted. The per-cell motif activities were computed by using the Signac wrapper for chromVAR [(*58*)](https://sciwheel.com/work/citation?ids=4187246&pre=&suf=&sa=0), which identifies motifs associated with variability in chromatin accessibility between cells.

#### Linking gene regulatory elements and gene expression

We examined the peak-gene relationships by using pseudobulk samples aggregating ATAC-seq and RNA-seq counts. We created these pseudobulk samples by randomly sampling 500 cells from the entire single cell dataset as seed cells. For each seed cell, we selected its 49 nearest neighbors within the cells of the same cell type annotations. Therefore, we obtained 500 pseudobulk samples, each of which comprised 50 single cells. The ATAC-seq and RNA-seq counts for pseudobulk samples were obtained by summing peak and gene counts across the respective single cell members.

We then evaluated the peak-gene linkage scores by using a correlation-based approach[(*14*, *21*, *31*)](https://sciwheel.com/work/citation?ids=9897011,10230238,5945818&pre=&pre=&pre=&suf=&suf=&suf=&sa=0,0,0) applied to these pseudobulk samples. We computed the Spearman correlation between all pairs of peaks and expression of nearby genes (within 0.5 Mbp) by using the LinkPeaks function implemented in Signac, and then selected all peak-gene pairs with correlation coefficient |ρ| > 0.3 and FDR-adjusted *P* value < 0.1. Additionally, we removed peak-gene links that overlapped the promoter region (defined as within -1000 to +100 bp from the TSS).

To define DORCs (a set of nearby peaks correlated with the expression of a target gene), we ranked genes by the number of significantly associated peaks among the peak-gene associations we identified above, and determined 5 peaks per gene as a cutoff based on the elbow method. To quantify DORC score for a gene, we used the sum of normalized counts in all significantly associated peaks per gene, resulting in a DORC x cell matrix.

Using the pseudobulk samples, we defined a normalized score (from 0 to 1) as the ‘pseudo-age’ for each cell type based on the proportion of cells found in the six different developmental stages. Taking into consideration the nonlinear developing speed across lifespan, we used log-scale weights, instead of equal weights, for each developmental stage: log_10_(1) = 0 for early fetal, log_10_(3) = 0.48 for late fetal, log_10_(5) = 0.70 for infancy, log_10_(7) = 0.85 for childhood, log_10_(9) = 0.95 for adolescence, and log_10_(10) = 1 for adulthood.

#### Pseudotime analysis on neuronal populations

We extracted cells which were annotated as neuronal cell types (including RG, IPC, EN-fetal-early, EN-fetal-late, EN, IN-fetal, IN-MGE and IN-CGE). We re-performed dimension reduction on the corresponding RNA-seq data and derived UMAP representations based on the top 10 PCs. The rationale behind using only the RNA-seq modality here is that RNA-seq provides higher resolution than ATAC-seq in terms of distinguishing cell types, especially maturely differentiated ones (**Fig. 1C**), which are usually the endpoints of trajectories and, hence, are necessary for accurate trajectory inference. There are 530 cells suspected as doublets of IN-MGE and IN-CGE (**fig. S3B**) and these were removed from the following pseudotime analysis. Next we used the UMAP coordinates as input to monocle3 [(*60*)](https://sciwheel.com/work/citation?ids=6482520&pre=&suf=&sa=0) to construct the trajectories across neuronal populations (parameters: learn_graph_control = list(minimal_branch_len = 20), use_partition = FALSE, close_loop = FALSE). To calculate pseudotime, we selected RGs as the ‘root’ nodes. For cell assignment to lineages, we excluded cells for which the cell type annotations were inconsistent. For example, any inhibitory neurons were excluded from the EN lineage, and any excitatory neurons were excluded from the IN-MGE or IN-CGE lineage. We used the graph_test function in monocle3 to find differentially expressed genes on the trajectory of a specific lineage, and selected genes with *q* value < 0.01.

For visualization and residual analysis purposes, we used the DORC scores and the associated gene expression, smoothed over the inferred pseudotime, by applying the tradeSeq package [(*66*)](https://sciwheel.com/work/citation?ids=8350898&pre=&suf=&sa=0) with nine knots. The ‘knots’ are points where a set of basis functions are joined together to create the smoothers. The number of knots was selected to reach an optimal bias-variance trade-off for the smoother, according to the Akaike information criterion (AIC) (**fig. S3B**). Specifically, we fit a negative binomial generalized additive model (NB-GAM) for every gene in each lineage by using the fitGAM function (parameter setting: ‘knots = 9’), and then obtain the estimated smoother by using the predictSmooth function. The residual for each gene was calculated by subtracting the min-max normalized gene expression values from the min-max normalized DORC scores.

#### Overlap with common genetic risk variants

To investigate whether the cell specific peaks and genes might play a role in various brain and non-brain related traits, we quantified their co-localization with common risk variants from 53 GWAS studies (**tables S10 and S11**). To perform this analysis for snATAC-seq data, we chose the top 2,500 most specific peaks per cell type (padded by 1 kb in both directions to include adjacent variants) and applied LD-score score partitioned heritability [(*39*)](https://sciwheel.com/work/citation?ids=5167004&pre=&suf=&sa=0). This method calculates if common genetic variants located within cell type specific peaks explain more of the heritability than variants not in those regions when adjusting for the number of variants and also for genetic context (e.g. introns, exons, promoters, or intergenic regions). To perform this analysis for snRNA-seq data, we selected the top 500 most specific genes per each cell type and applied MAGMA [(*38*)](https://sciwheel.com/work/citation?ids=1234481&pre=&suf=&sa=0). Cell specific genes were padded by 35 kb upstream and 10kb downstream to also include genetic variants in the proximal regulatory regions. For all GWAS studies, we excluded the broad MHC-region (hg19 coordinates on chr6: 25-35Mb) due to its extensive and complex LD structure. Linkage disequilibrium (LD) was estimated from the European panel of 1000 Genome Project phase 3. For the AD GWAS, we removed the APOE effect in the model by excluding SNPs around the APOE gene. For MDD GWAS, we used versions of summary statistics without 23andMe individuals. Summary statistics were downloaded from the designated locations mentioned in the original manuscripts that are referred to in **tables S10 and S11**.

To link risk loci associated with neuropsychiatric disorders to their causal genes, we first collected a list of genome-wide significant variants of those disorders extended by the variants that are in strong LD (R≥0.8). For this purpose, we utilized LD matrix for 1000 Genomes Phase 1 of individuals of European ancestry downloaded from https://zenodo.org/record/3404275#.YaQec5HMJdC. Then, we overlapped these variants with the peaks for which we have at least one peak-gene defining the gene under regulation (peaks were lifted to hg19 to match default hg19 coordinates of index SNPs and LD buddies).

#### Validation Experiments

**Guide Design**

The CRISPRi algorithm of the Broad Institute Genetic Perturbation Platform was used to design guide RNAs against regions proximal to the target gene TSS (transcription start site).

| **Guide Name** | **Sequence** | **PAM** | **sgRNA target Chromosomal position** |
| --- | --- | --- | --- |
| NEUROD1 1 | AGTGATAGTCTCATAACCCT | GGG | Chr2: 181680471 |
| NEUROD1 2 | TTATGAGACTATCACTGCTC | AGG | Chr2: 181680453 |
| NEUROD1 3 | GCAGGAGGCGCGGCGTCCGG | AGG | Chr2: 181680484 |
| CUX2 1 | GAGATGCAGCGAGCGCTCCG | CGG | Chr12:111033777 |
| CUX2 2 | AGATGCAGCGAGCGCTCCGC | GGG | Chr12:111033778 |
| CUX2 3 | AGCGAGCGCTCCGCGGGCCC | GGG | Chr12:111033784 |
| Scrambled | GCACTCACATCGCTACATCA |  |  |

**Plasmid Preparation**

The top 3 guide sequences for each target and their reverse complement sequences were synthesized by IDT (Integrated DNA technologies) with the appropriate overhang sequences to be cloned downstream of a constitutively expressed U6 promoter in a lentiviral vector (lentiGuide-Hygro-mTagBFP2, Addgene, Cat# 99374) using the golden gate cloning method. Guide oligos and their reverse complement oligos were initially phosphorylated and annealed using the following protocol:

Reaction Mix:

1 μl guide oligo (100 μM)

1 μl reverse complement oligo (100 μM)

1 μl 10X T4 DNA Ligase buffer (New England Biolabs, Cat# B0202S)

0.5 μl T4 PNK (New England Biolabs, Cat# M0201L)

6.5 μl ddH2O

10 μl total

Thermocycler:

37℃ 30 min

95℃ 5 min

Ramp down to 25℃ at 5℃ /min

4℃ infinite hold

Phospho-annealed oligos were diluted 1:100 before proceeding to the golden gate reaction:

Reaction Mix:

12.5 μl Quick Ligation Buffer (2X) (New England Biolabs, Cat# M2200L)

0.25 μl BSA (10 mg/ml)

1 μl BsmB1 v2 (New England Biolabs, Cat# R0739L) (optimized for golden gate cloning)

0.125 μl T7 DNA Ligase (New England Biolabs, Cat# M0318S)

1 μl diluted phospho-annealed oligos

1 μl of [25ng/ul] lentiGuide-Hygro-mTagBFP2

9.125 μl ddH_2_O

25 μl total

Thermocycler:

Cycle (30x):

37℃ 5 min

20℃ 5 min

4℃ infinite hold

Ligations were transformed into High Efficiency NEB 10-beta Competent *E. coli*. (New England Biolabs, Cat# C3019H) according to manufacturer’s instructions. Bacteria were plated on LB-Ampicillin agar medium at 37℃ overnight. On the following day, colonies were grown in LB-Ampicillin liquid medium at 37℃ with shaking. Plasmids were purified with the QIAprep Spin Miniprep kit (Qiagen, Cat# 27106). Positive clones were validated through sanger sequencing by GeneWiz using a universal U6 forward primer. Validated plasmid DNA containing guide RNA inserts were packaged into lentivirus by VectorBuilder with a viral titer of >10^8^ TU/ml in HBSS Buffer.

**Transduction /Differentiation**

hiPSC-NPCs were differentiated into forebrain neurons as previously described [(*36*)](https://sciwheel.com/work/citation?ids=7889160&pre=&suf=&sa=0). hiPSC-NPCs were seeded at low density with 1uL of lentivirus per mL of NPC media (DMEM/F12, 1x N2, 1x B27- RA (Invitrogen), 1 μg/ml laminin and 20 ng/ml FGF2 in Matrigel-coated plates. 1-2 days after plating cells were treated with 1 mg/mL hygromycin (ThermoFisher Scientific) to select for cells containing the gRNA. 2 days post selection, cells were cultured in neural differentiation medium (DMEM/F12 + Glutamax, 1x N2, 1x B27-RA, 20 ng/ml BDNF (Peprotech), 20 ng/ml GDNF (Peprotech), 1 mM dibutyryl-cyclic AMP (Sigma), 200 nM ascorbic acid (Sigma) and 1 μg/ml laminin (ThermoFisher Scientific)). hiPSC-derived-(forebrain)-neurons were differentiated for 2-6 weeks depending on the assay.

**RNAscope setup (Plating)**

hiPSC-NPCs were plated into 8-well chamber slides (Lab-Tek) at a density of 3.0 x 10^4^ per well for RNAscope. Lentiviral transduction and differentiation was carried out as previously described. Cells were fixed for RNAscope at 2 weeks post differentiation.

**RNAscope**

Growth media was removed from the 8-well chamber slides and cells were washed with 1x PBS (500 μl per chamber). Cells were then fixed with 4% formaldehyde/1x PBS for 30 min at RT (500 μl per chamber). After fixation, cells were washed 3x with 1x PBS and dehydrated with 50%, 70%, and then 100% EtOH according to instructions from Advanced Cell Diagnostics for adherent cells grown in chamber slides. Chambers were removed and the slides were stored in 100% EtOH at -20℃. Cells were rehydrated and RNAscope was performed according to manufacturer’s instructions. Primary probes targeting NEUROD1 (Advanced Cell Diagnostics RNAScope Probe- Hs-NEUROD1-C2 (#437281-C2)) were hybridized and amplified with secondary and tertiary probes, which were then labeled with the fluorophore Opal 570 (Akoya Biosciences FP1488001KT). Primary probes targeting CUX2 (Advanced Cell Diagnostics RNAscope Probe- Hs-CUX2-C3 (#425581-C3) were hybridized and amplified with secondary and tertiary probes, which were then labeled with the fluorophore Opal 690 (Akoya Biosciences FP1497001KT). Cells were counterstained with DAPI and mounted with Prolong Gold Antifade Reagent (Thermo Fisher P36934). Mounting media was allowed to dry overnight and then slides were stored in the dark at 4 C before cells were imaged.

RNAscope slides were imaged with the Zeiss Axioimager.Z2(M) widefield microscope with a Axiocam monochrome 503 CCD camera (Pixel size: 4.54um X 4.54um). A 10x air/dry objective (Fluar 10x/NA 0.5) was initially used to take snapshots of large fields of cells (400 ms exposure), and then a 63x oil immersion (PlanApo 63x/NA 1.4) objective was used to capture images at higher magnification for further quantitative analysis. With the 63x oil objective, 35-40 z stacks (each z stack is 200 nm) were acquired in 3 channels: Cy5 (Chroma 49006 filter cube) to image CUX2, Cy3 (Chroma 49309 filter cube) to image NEUROD1, and DAPI (Chroma 49000 filter cube). For Cy5 and Cy3, the exposure time for each z stack was 200 ms, and for DAPI it was 50 ms.

Maximum intensity projections were generated from the Cy5 and Cy3 z stacks using FiJi. CUX2 (Cy5) and NEUROD1 (Cy3) RNAscope spots in the maximum intensity projections were localized using the IDL script LocalizeApp, which utilizes 2D gaussian fitting to calculate the center of diffraction limited spots. For the DAPI z stack, the z plane in which nuclei were in focus was used for manual segmentation of nuclei and generation of binary nuclear masks. The IDL script FISHauxiliary was used to quantify the number of diffraction limited spots in each nuclear mask for both CUX2 and NEUROD1.

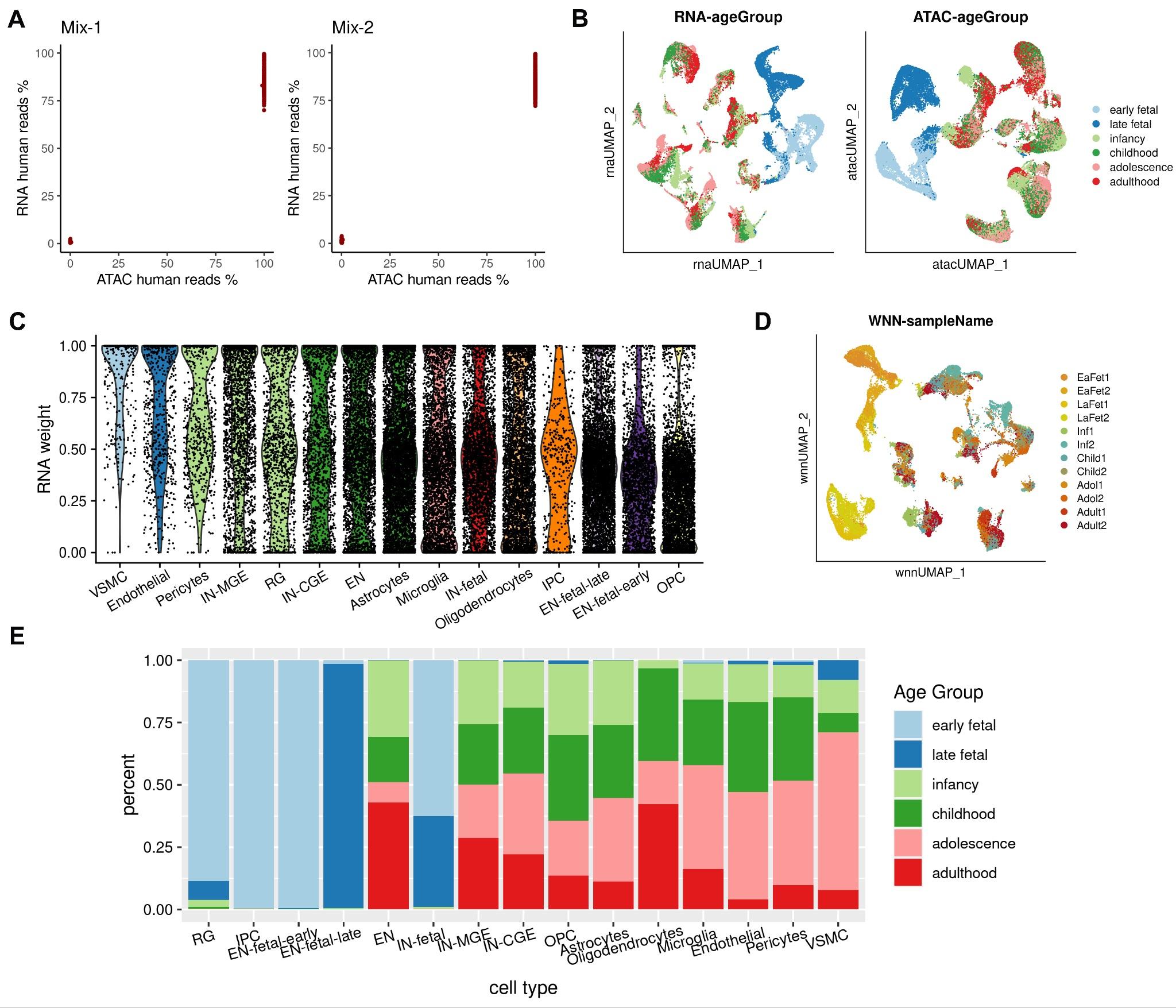

**Fig. S1.**

**(A)** The percentage of ATAC-seq or RNA-seq reads aligning to the human genome relative to all reads mapping uniquely to the human or mouse genome in the human-mouse cell line mixtures. We retained cells with total RNA-seq count > 2,000 and < 9,500, total ATAC-seq count > 15,000 and < 200,000, and mitochondrial percentage < 30%. The cell numbers for the two mixed samples are: 465 human cells and 319 mouse cells in Mix-1, 439 human cells and 312 mouse cells in Mix-2. **(B)** UMAP visualizations of single cells defined by RNA-seq and ATAC-seq data, respectively, where cells are annotated in terms of the age groups. **(C)** UMAP visualization of single cells based on the WNN-derived graph, where cells are annotated in terms of sample names. **(D)** Single cell RNA modality weights derived from WNN analysis. VSMCs, endothelial cells and pericytes have the highest RNA weights, consistent with the fact that they can be distinguished only in RNA modality. **(E)** Bar plots showing the proportion of cells coming from different age groups for each cell type.

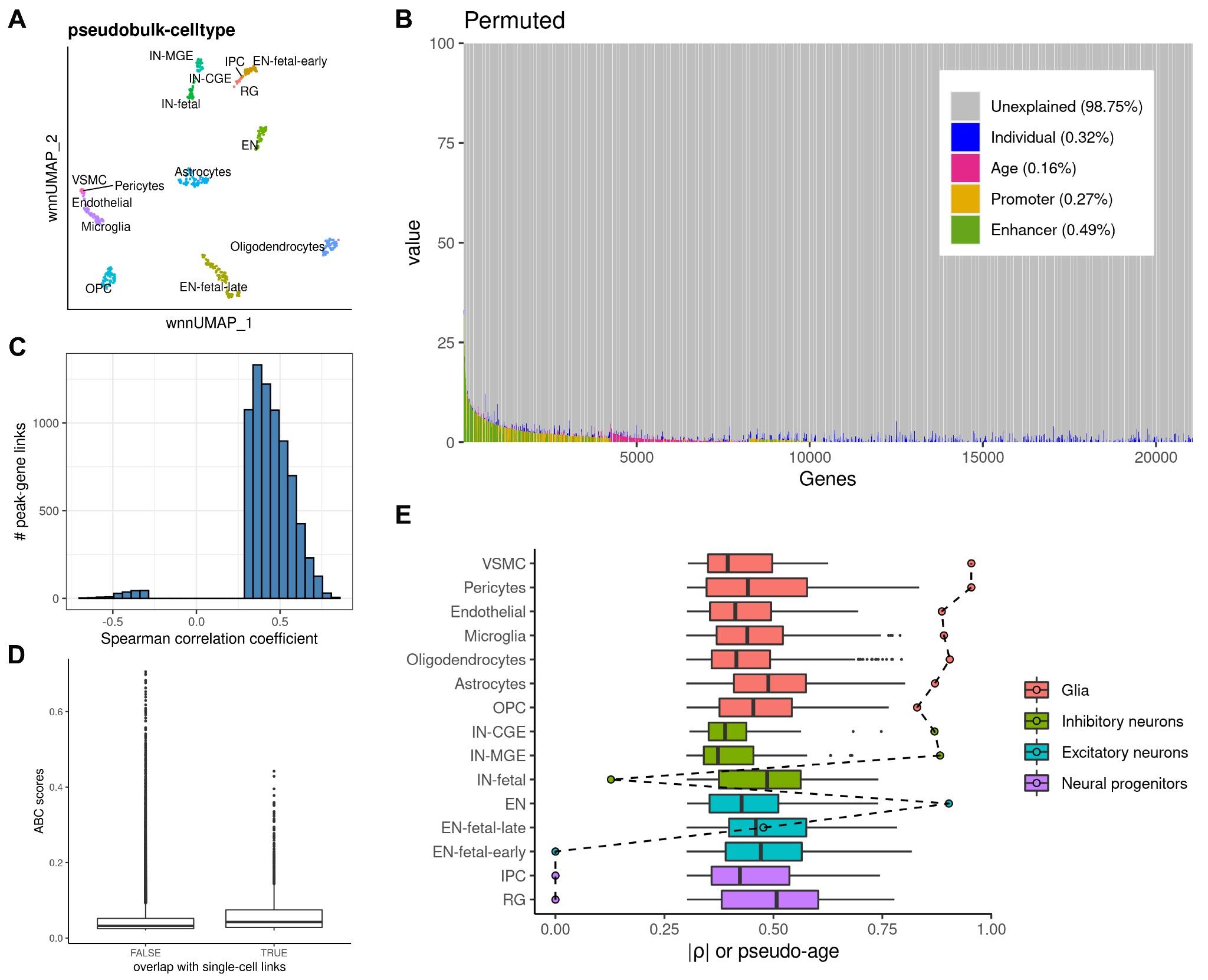

**Fig. S2.**

**(A)** WNN-derived UMAP visualization for the 500 pseudobulk aggregate samples, annotated by cell types. **(B)** Variance in gene expression estimated to be explained by chromatin accessibility, as well as other variables, using shuffled data. Genes, in columns, are sorted by decreasing proportion of variance explained by the epigenome (enhancers and promoters), with the mean variance explained by each component shown in parenthesis. **(C)** Histograms showing the Spearman correlation coefficients of the significant peak-gene links. **(D)** Boxplots for comparing the ABC scores of the E-P interactions overlapping peak-gene links, versus those not overlapping. **(E)** Boxplots showing the distribution of the absolute Spearman correlation coefficients of the peak-gene links specific to different cell types; circles connected by a dashed line represent the pseudo-age per cell type; colored in terms of broader cell type classification.

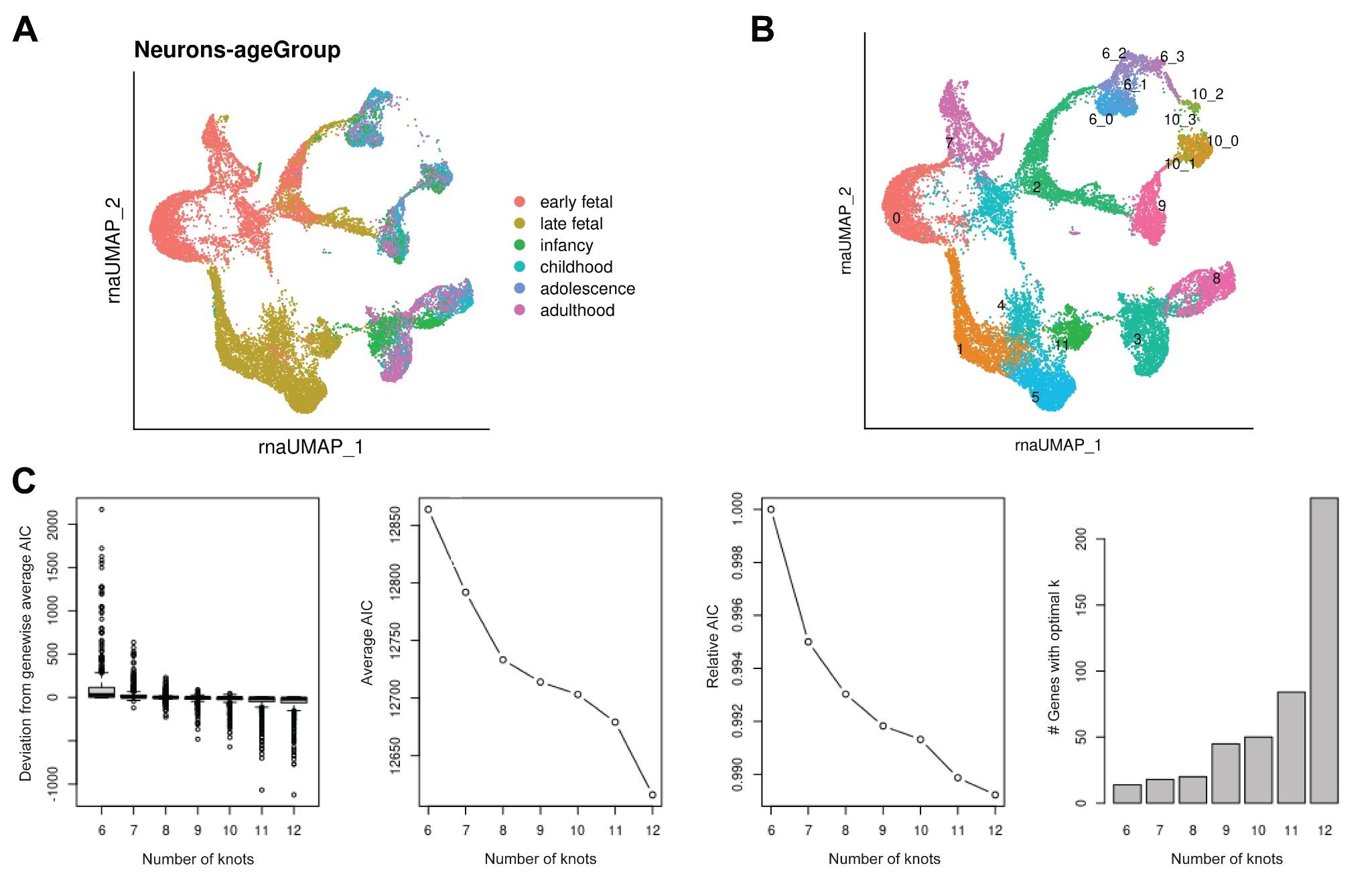

**Fig. S3.**

**(A)** UMAP visualization of neuronal cell populations on RNA coordinates, colored in terms of different developmental stages. **(B)** UMAP visualization of neuronal cell populations, colored and labeled in the cluster IDs. Cells assigned to clusters ‘6_3’ (n=380) and ‘10_2’ (n=150) were suspected as doublets between IN-MGE and IN-CGE, and were removed from downstream pseudotime analysis. **(C)** Diagnostic plots for selecting the optimal number of knots, k ∈ {6, …, 12}, using the AIC as implemented in the evaluateK function in tradeSeq [(*66*)](https://sciwheel.com/work/citation?ids=8350898&pre=&suf=&sa=0). See tradeSeq tutorials for more details.

**Table S1.**

Sample information including brain regions, age, sex, and batches.

**Table S2.**

Differentially expressed genes for each cell type.

**Table S3.**

Differentially accessible peaks for each cell type.

**Table S4.**

List of peak-gene links which were identified with significant associations.

**Table S5.**

List of glia-/neuron-specific super-enhancers overlapped with each DORC.

**Table S6.**

Differentially expressed DORC-regulated genes for each cell type.

**Table S7.**

List of peak-gene links which were identified among neurons with significant associations.

**Table S8.**

GO enrichment analysis results for neuron-specific DORC-regulated genes.

**Table S9.**

TF motif enrichment results for neuronal lineage-specific clusters of genes (km1/2/3/4).

**Table S10.**

Overlap between common risk genetic variants associated with 53 brain and non-brain related traits and cell-specific peaks as calculated by LD-sc method.

**Table S11**.

Overlap between common risk genetic variants associated with 53 brain and non-brain related traits and marker genes (including proximity of those genes) as calculated by MAGMA method.

**Table S12**.

Candidate causal genes for risk variants (GWAS index SNP and their LD buddies) associated with neuropsychiatric disorders.
